# Enhanced lentiviral gene delivery to mammalian cells via paired cell surface and viral envelope engineering

**DOI:** 10.1101/2024.11.08.622723

**Authors:** Ryan Tannir, Letitia Chan, Tomasz M. Grzywa, Orlando Arevalo, Alexandra Neeser, Sara Kahn, Austin Cozzone, Lauren Olenick, Nicola J Mason, Leyuan Ma

## Abstract

Lentiviral vectors that facilitate gene delivery to desired cell types have been widely used in routine laboratory research and therapeutic cell engineering. However, the lack of proper entry receptors on many cell types often results in poor gene delivery. Here, we present a simple paired virus-cell engineering approach that promotes lentiviral gene delivery into mammalian cells. Lentiviruses are dual-pseudotyped with VSV-G and a chimeric envelope protein specifically recognizing a small molecule fluorescein (αFITC-Env), and target cells are transiently labelled with FITC to create surrogate receptors for lentivirus attachment. The synthetic interaction between FITC-labeled cells and FITC-binding LVs enables efficient LV docking, viral entry and stable transgene expression in a range of mammalian cell lines and primary T cells. We showed that this approach enabled efficient delivery of a CD19-targeted chimeric antigen receptor (CAR) into naïve human T cells that are naturally refractory to conventional VSV-G LVs, which upon activation rapidly eradicated CD19^+^ leukemic cells. This paired cell surface and virus envelope engineering approach may serve as a universal method for engineering synthetic virus-cell interactions to improve lentiviral gene delivery to mammalian cells.

## INTRODUCTION

Lentiviral vectors (LVs) are derived from human immunodeficiency virus type 1 (HIV-1) and are replication-incompetent for enhanced safety^1^. LVs have become indispensable tools in preclinical research and therapeutic cell engineering due to their ability to stably integrate genetic materials into both dividing and non-dividing mammalian cells^2^. Notably, LVs are the primary gene delivery vector for the generation of Chimeric Antigen Receptor (CAR) T cells or T cell receptor (TCR)-transgenic T cells^3^. Central to the efficacy of LVs is lentiviral pseudotyping, which involves incorporating envelope proteins (Env) from alternative non-HIV viral sources to change the LV tropism^4^. For example, RabV-G from Rabies virus was used for targeting neurons^5^, H/F from measles virus for targeting B and T cells^6^, and G/F from Nipah virus for targeting pericytes and tumor endothelium^7^. However, VSV-G remains widely used owing to its broad tropism, stability, as well as the high transduction efficiency of VSV-G pseudotyped LVs^8^. VSV-G predominantly interacts with low-density lipoprotein receptors (LDLR) and possibly other LDLR family receptors present on the cell surface^9^. Binding of VSV-G to the LDLR triggers clathrin-mediated endocytosis, a pH-mediated membrane fusion, and subsequent viral genome delivery into target cells^10^.

Various approaches have been leveraged to improve LV transduction. For example, a cationic polymer, polybrene, that neutralizes repulsive charges between the virus and the cell membrane, could facilitate viral attachment and entry into target cells^11^. Retronectin, a recombinant protein that binds to VLA-4 and VLA-5 integrins on the cell surface, could bring the virus closer to the cell surface to promote viral attachment and entry^12^. However, many cell types remain refractory to VSV-G LV transduction with polybrene or retronectin, including hematopoetic stem cells in G0 phase^13^, monocytes^14^, quiescent T^15^ and B cells^16^, as well as murine T cells^17^ and canine T cells^18^. While this refraction could be due to a number of mechanisms, such as poor viral entry due to a lack of LDLR^19^, lack of ATP-dependent nuclear import^20^, poor reverse transcription^21^, or poor integration^22^, efficient of LV binding to a proper entry receptor seems to be deterministic of LV transduction in most occasions^23^. These were exemplified by pseudotyping LVs with chimeric Envs with single-chain variable fragment (scFv)-directed toward specific surface proteins, such as anti-CD4/CD8 scFv pseudotyping for transducing T cells^24^. However, finding proper entry receptors to achieve efficient LV gene delivery for each cell type, especially primary cells, could be a daunting task.

Here, we demonstrate a surface chemistry-assisted LV transduction as a generic approach to enhance LV gene delivery to mammalian cells (**Fig. 1A**). Chemically attaching fluorescein (FITC) to cell surfaces coupled with VSV-G and αFITC-Env (αFITC-Env) dual-pseudotyped LVs provide tailored virus binding, entry, and enhanced LV gene delivery to many types of mammalian cells, including naïve human T cells, for therapeutic T cell engineering.

**Figure 1:**
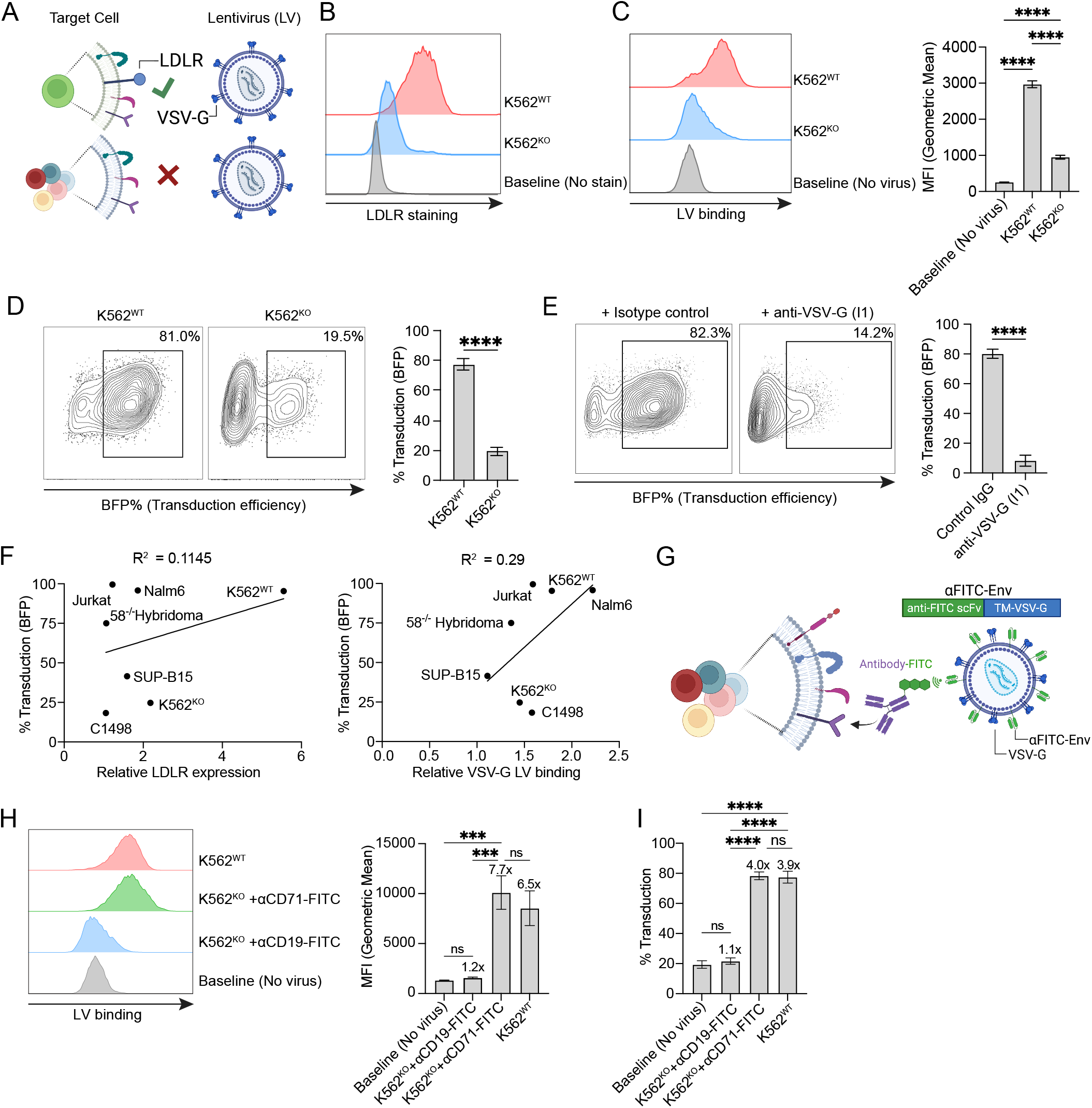
Active virus-cell interactions are required for successful lentiviral transduction. **(A)** Schematic representation of receptor requirement for lentiviral attachment and transduction. **(B)** Histogram plot of human LDLR level on K562^WT^ and K562^KO^ cells detected by a PE-conjugated anti-LDLR antibody using flow cytometry. **(C)** VSV-G virus binding to K562^WT^ and K562^KO^ cells detected using anti-VSV-G antibody and flow cytometry. Shown are a representative histogram and summary results of the The mean fluorescence intensity (MFI). **(D)** VSV-G LV transduction (BFP^+^) of K562^WT^ and K562^KO^ cell. Shown are the representative scatter plots and summary results. **(E)** Transduction of K562^WT^ cells using VSV-G LV pre-blocked with an anti-VSV-G antibody (clone: I1). Shown are the representative scatter plots and summary results. **(F)** The correlation between relative LDLR expression or relative VSV-G virus binding and LV transduction of different mammalian cell lines. **(G)** Schematics showing VSV-G/αFITC-Env dual-pseudotyped LV. **(H-I)** αFITC/VSV-G LV binding to (**H**) and transduction (**I**) of K562^KO^ cells pre-labeled with αCD71-FITC or control αCD19-FITC. K562^WT^ cells were included as control. Shown are representative histogram and summary results of the MFI for H and transduction% for I. Fold increase above the baseline was shown for each condition. Throughout this figure, BFP was used as a reporter and transduction efficiently was determined as the BFP level 48 hours post transduction. Error bars show mean ± SD (n=3). **, P<0.01; ***, P<0.001; ****, P<0.0001 by one-way ANOVA with Tukey’s post-test for C, H and I; by unpaired t-test for D and E. For F, curve fitted and R^2^ calculated by simple linear regression.

## RESULTS

### Active natural and synthetic virus-cell interaction is required for successful LV transduction of mammalian cells

To investigate the receptor dependence of VSV-G LV gene delivery, we focused on LDLR+ K562 (K562^WT^) cells and generated an isogenic LDLR-deficient K562 cell line using CRISPR Cas9 editing, termed LDLR-KO K562 (K562^KO^) (**Fig. 1B**), which exhibited significantly reduced VSV-G LV binding (**Fig. 1C)**. Using a Blue Fluorescent Protein (BFP) as a reporter gene, we showed that K562^KO^ cells have a marked reduction of VSV-G LV transduction from >80% to <20% (**Fig. 1D**), analogous to abolished transduction of K562^WT^ cells by blocking LDLR-VSV-G interaction using an anti-VSV-G antibody (Clone: I1) (**Fig.1E**). These results are consistent with the notion that LDLR is the natural entry receptor for VSV-G LV.

Different cell types have different expression levels of LDLR and potentially other suboptimal VSV-G LV binding receptors^9^. We next sought to determine the role of LDLR versus overall virus binding in VSV-G LV transduction. To this end, we selected a panel of different mammalian cells, namely, Nalm6, Jurkat, Raji, 58^-/-^ hybridoma, C1498, SUP-B15, and performed flow cytometry analysis for LDLR expression and VSV-G LV binding. Despite that all cell lines possess detectable LDLR expression, LV binding to the cell surface exhibited a stronger correlation with the ultimate transduction efficiency than LDLR expression (R^2^= 0.29 vs 0.11) (**Fig. 1F**).

Finally, to assess if synthetic LV binding could facilitate LV transduction, we designed a chimeric envelope protein by replacing the extracellular domain of VSV-G with a high-affinity anti-FITC single-chain variable fragment (αFITC-Env, Clone E2) and generated LVs bearing both VSV-G and αFITC-Env (αFITC/VSV-G LV hereafter) (**Fig.1G**). K562^KO^ cells were pre-labeled with a FITC-conjugated CD71-targeted antibody (αCD71-FITC) to allow FITC as a surrogate surface receptor for αFITC/VSV-G LV (**Fig. 1H**), and this synthetic interaction enabled nearly complete restoration of LV transduction (**Fig. 1I**). These results suggest that efficient VSV-G LV transduction requires active virus-target cell interaction, which could be achieved by either natural or synthetic binding mechanisms.

### Passive virus attachment onto cell surface supports but is not sufficient for optimal LV transduction

Physically increasing the adjacency of LV particles to target cells is critical for efficient LV transduction^25^. The positively charged polymer polybrene promotes LV transduction by enabling the co-localization of virus and cells via surface charge neutralization^10^. To test if polybrene is enough to enable VSV-G LV localization to cell surface, subsequent virus entry and stable gene delivery, K562^WT^ and K562^KO^ cells were labeled with VSV-G LV in the presence or absence of polybrene. Polybrene significantly increased VSV-G LV attachment to both K562^WT^ and K562^KO^ cells, with stronger attachment to K562^WT^ cells (**Fig. 2A**). However, strong virus attachment to K562^KO^ cell surface only led to a mild increase of LV transduction from around 20% to 40% (**Fig. 2B**), likely through a receptor-independent virus entry mechanism^26^. Polybrene also improved αFITC/VSV-G LV transduction of αCD71-FITC pre-labeled K562^KO^ cells (**Fig. 2C**). Notably, blocking VSV-G did not prevent VSV-G LV attachment to cell surface in the presence of polybrene (**Fig. 2D**) but prevented transduction of K562^WT^ cells (**Fig. 2E**). Additionally, the use of a different LV transduction enhancer, retronectin, which binds both viruses and integrins on the target cell surface to bring viruses closer to the cell^11^, failed to improve LV transduction of K562^KO^ cells **(Fig. 2F)**. These results suggest that active virus binding to the cell surface via Env-receptor interaction plays a determining role in LV transduction, which could be further improved by non-specific attachment or physical proximity of the virus particles to the cell surface.

**Figure 2:**
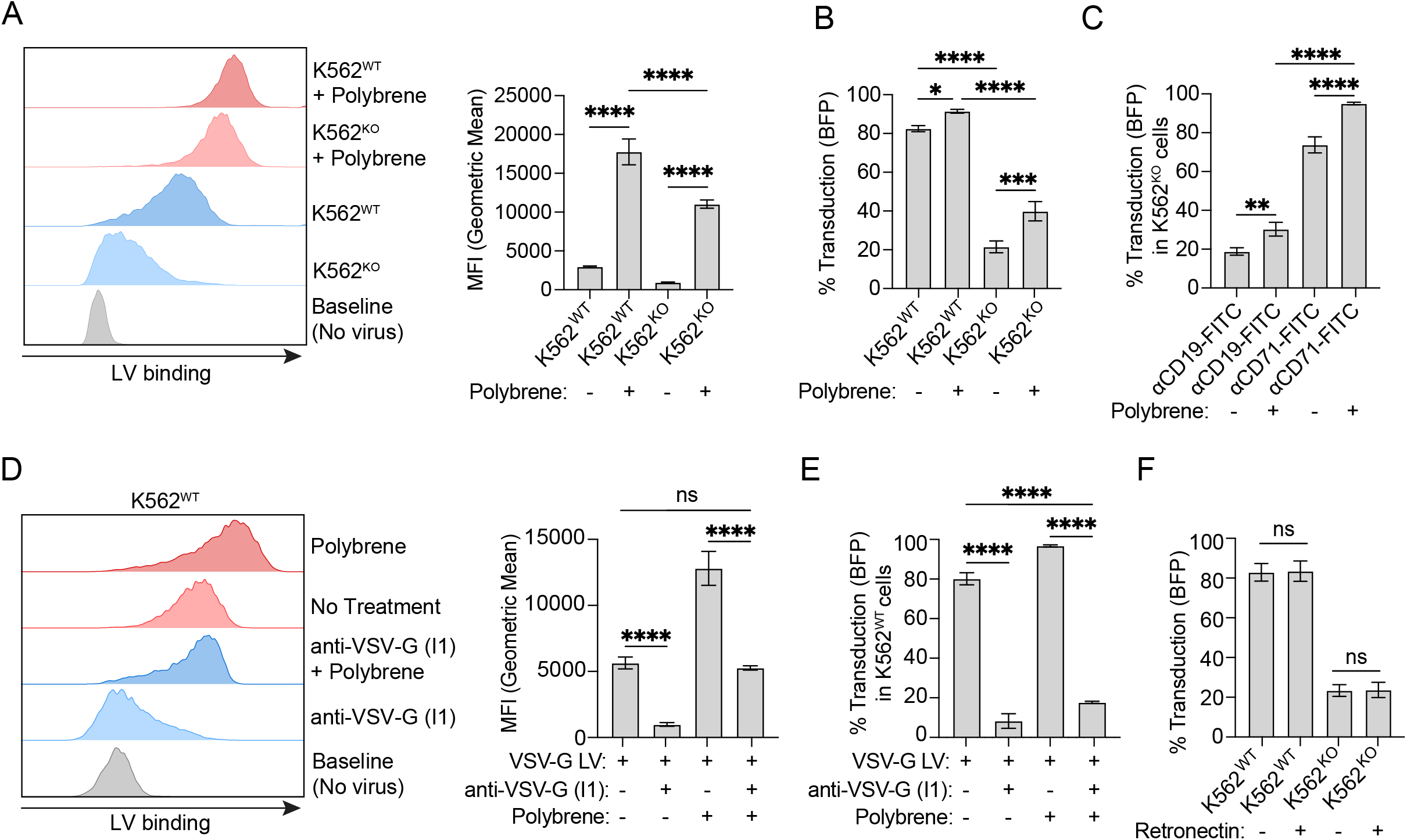
Virus localization to cell surface is required but not sufficient for lentivirus transduction. **(A-B)** VSV-G LV binding to (**A**) and transduction (**B**) of K562^WT^ and K562^KO^ cells with or without polybrene. Shown are representative histogram and summary results of the MFI for A and transduction% for B. **(C)** αFITC/VSV-G LV transduction of K562^KO^ cells pre-labeled with αCD71-FITC or control αCD19-FITC with or without polybrene. (**D-E)** Impact of VSV-G blockade on VSV-G LV binding to (**D**) and transduction (**E**) of K562^WT^ and K562^KO^ in the presence or absence of polybrene. Shown are representative histogram and summary results of the MFI for D and transduction% for E. **(F)** Transduction of K562^WT^ and K562^KO^ cells with VSV-G LV in the presence or absence of retronectin. Throughout this figure, BFP was used as a reporter and transduction efficiently was determined as the BFP level 48 hours post transduction. Error bars show mean ± SD (n=3). *, P<0.05; **, P<0.01; ***, P<0.001; ****, P<0.0001; ns, non-significant by one-way ANOVA with Tukey’s post-test for A-F.

### Chemical modification of cell surface bypasses the natural entry receptor to promote LV transduction

Building on our observation that αCD71-FITC mediated cell surface labeling could restore LV transduction, we reason that non-selective chemical attachment of FITC to target cell surfaces could be a universal approach for improving LV transduction. We recently found that an amphiphilic lipid conjugate, DSPE-PEG2k-FITC (amph-FITC^2k^), could efficiently insert into the lipid bilayer of any target cell surface^27^. We sought to determine if the specific and high-affinity attachment of αFITC/VSV-G LV to cell surface through amph-FITC could promote LV transduction (**Fig. 3A**). Labeling of K562^KO^ cells with amph-FITC^2k^ demonstrated a strong increase in αFITC/VSV-G LV binding, comparable to the binding to K562^WT^ (**Fig. 3B**). Accordingly, only after amph-FITC^2k^ labeling, K562^KO^ cells were transduced efficiently with αFITC/VSV-G LV (**Fig. 3C**). We next sought to explore how the length of PEG linker within amph-FITC affects transduction. Labeling K562^KO^ cells with amph-FITC bearing various linker lengths confirmed that a 5k PEG linker led to an optimal 3.8 folds increase of αFITC/VSV-G LV transduction (**Fig. 3D**) compared to the baseline (labeling with free FITC).

**Figure 3:**
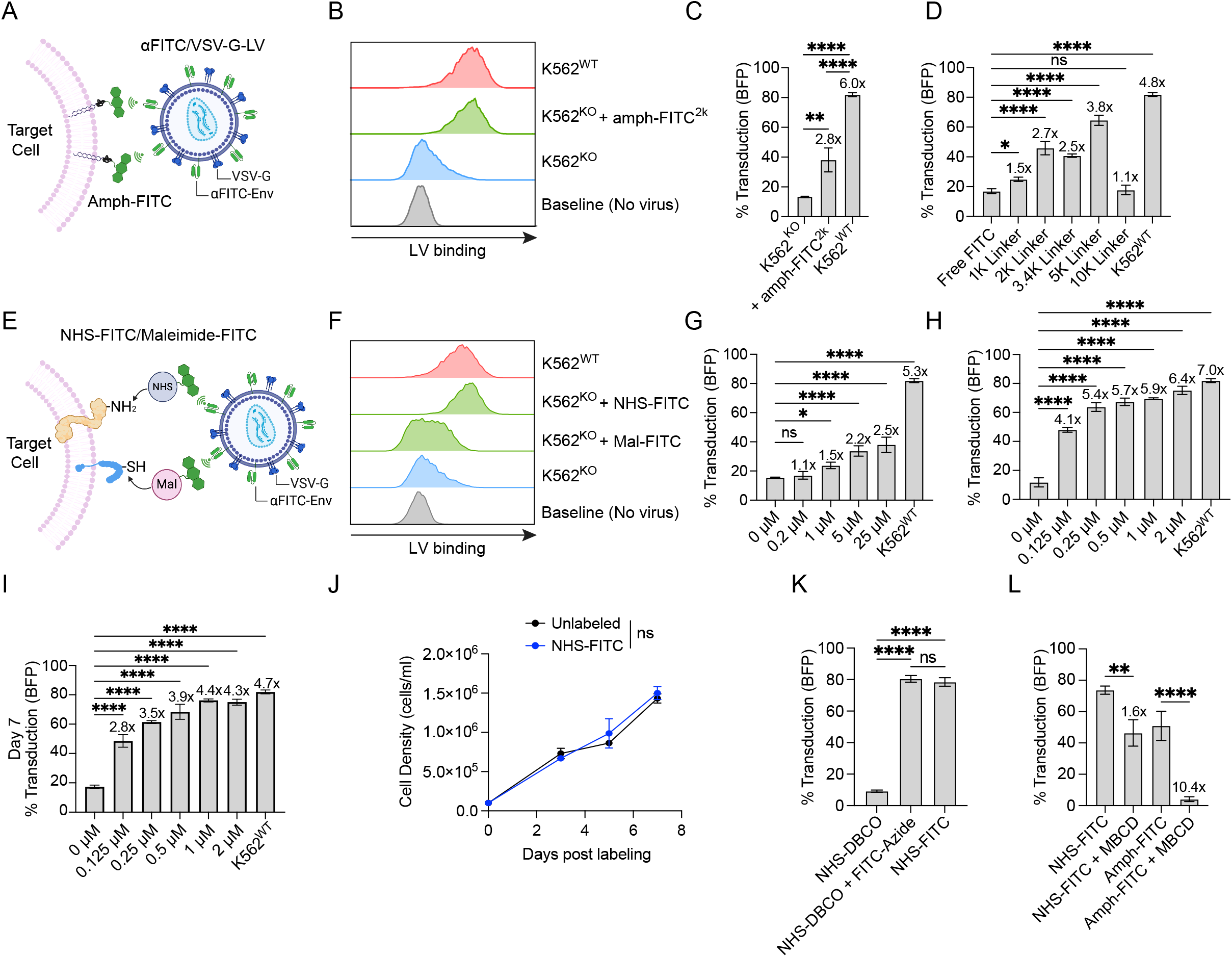
Chemical modification of the cell surface bypasses the natural entry receptor to enable LV transduction. **(A)** Schematic representation of αFITC/VSV-G LV transduction of amph-FITC labeled cells. **(B)** Representative histograms of αFITC/VSV-G LV binding to K562^KO^ cells labeled with amph-FITC^2k^ (1μM) or amph-control. **(C)** αFITC/VSV-G LV transduction of K562^KO^ cells pre-labeled with or without amph-FITC^2k^ (1μM). **(D)** αFITC/VSV-G LV transduction of K562^KO^ cells pre-labeled with free FITC or amph-FITC (1μM) with indicated PEG linker lengths. **(E)** Schematic representation of αFITC/VSV-G LV transduction of NHS-FITC or Mal-FITC label cells. **(F)** Representative histograms of αFITC/VSV-G LV binding to K562^KO^ cells labeled with either Mal-FITC (5 μM) or NHS-FITC (1 μM). **(G)** αFITC/VSV-G LV transduction of K562^KO^ cells pre-labeled with Mal-FITC at indicated concentrations. K562^WT^ shown as a control. **(H)** αFITC/VSV-G LV transduction of K562^KO^ cells pre-labeled with NHS-FITC at indicated concentrations. **(I)** Stability of αFITC/VSV-G LV transduction of NHS-FITC (1μM) labeled K562^KO^ cells 7 days post transduction. **(J)** Viability of NHS-FITC(1μM)-labeled K562^KO^ cells following transduction. Unlabeled K562^KO^ cells shown as control. (**K)** Schematics showing two-step labeling of cells with NHS-DBCO and FITC-Azide for αFITC/VSV-G LV transduction. **(L)** αFITC/VSV-G LV transduction of K562^KO^ cells labeled via NHS-DBCO (1μM) +/-FITC-Azide(1μM) or NHS-FITC (1μM). Throughout the figure, K562^WT^ cells were included as control, and BFP was used as a reporter and transduction efficiently was determined as the BFP level 48 hours post transduction. Fold increase above the baseline was shown for each condition in C, D, G and H. Error bars show mean ± SD (n=3). *, P<0.05; **, P<0.01; ***, P<0.001; ****, P<0.0001; ns, non-significant by one-way ANOVA with Tukey’s post-test for C and K; with Dunnet’s post-test for D, G, H, I; with Šidák’s post-test for L; by two-way ANOVA with Tukey’s post-test for J.

In addition to cell surface labeling with amph-FITC via direct lipid insertion, mammalian cell surface proteins bear various bio-active functional groups such as the free thiols and primary amines that can be leveraged for incorporating FITC via a covalent chemical conjugation (**Fig. 3E**). To this end, we labeled K562^KO^ cells with maleimide-FITC (Mal-FITC) or N-Hydroxysuccinimide-FITC (NHS-FITC) to covalently attach FITC onto the free thiol or primary amine groups on the cell surface, respectively. While Mal-FITC labeling of K562^KO^ cells only slightly increased αFITC/VSV-G LV binding, NHS-FITC labeling completely restored αFITC/VSV-G LV binding to K562^KO^ cells to the wildtype level (**Fig. 3F**). Accordingly, Mal-FITC labeling moderately improved the transduction of K562^KO^ cells from less than 20% to around 40% (**Fig. 3G**), and NHS-FITC labeling greatly increased the transduction to more than 70%, comparable to the transduction of unmodified K562^WT^ cells (**Fig. 3H**). Continuously monitoring transgene expression for 7 days confirmed stable transgene expression, thus ruling out the possibility of pseudo-transduction (**Fig. 3I**). Notably, these surface modifications have negligible impact on cell viability and proliferation (**Fig. 3J**). To rule out the possibility of NHS chemistry itself being a major contributor, we performed a two-step labeling. K562^KO^ cells were first labeled with Dibenzocyclooctyne-N-hydroxysuccinimidyl ester (NHS-DBCO), followed by covalent attachment of FITC-Azide via DBCO-Azide click chemistry (**Fig. 3K**). While NHS-DBCO labeling itself does not improve LV transduction, the subsequent addition of FITC-Azide led to a comparable LV transduction as direct NHS-FITC labeling (**Fig. 3L**).

### Paired cell surface FITC labeling and αFITC-Env pseudotyping as a generic approach to improve LV transduction of established mammalian cell lines and primary canine T cells

Given the orthogonality of each labeling chemistry and their independent mechanisms in promoting LV transduction, we sought to determine if a combination of these labeling methods could further improve the transduction efficiency of mammalian cells. Pre-labeling K562^KO^ cells with a cocktail of amph-FITC^5k^, Maleimide-FITC and NHS-FITC at the optimal condition yielded a comparable or worse transduction efficiency compared to NHS-FITC alone, highlighting the dominating effect of NHS-FITC in promoting LV transduction (**Fig. 4A**). Therefore, we focused on NHS-FITC solo labeling for all future LV transductions. Using a panel of mammalian cells, we found that NHS-FITC labeling significantly increased αFITC/VSV-G LV transduction of these cell lines by 1.4 – 54 folds (**Fig. 4B**). Using this platform, we also successfully integrated a membrane-bound human CD19 onto the surface of K562^KO^ cells and this expression was further increased with the addition of polybrene **(Fig. 4C)**, showing that cargo delivery is not limited to intracellular BFP. Next, we sought to determine if this surface FITC labeling-assisted LV transduction approach could improve the gene delivery to primary canine T cells, which are known to be resistant to VSV-G LV transduction^18^. Consistent with the earlier report, standard VSV-G LV transduction of activated canine T cells in the presence of polybrene led to minimal transduction (**Fig. 4D-E**), yet NHS-FITC labeling followed by transduction with αFITC/VSV-G LV markedly increased gene delivery to canine T cells from 1.63% to 40.3% (**Fig. 4D-E**).

**Figure 4:**
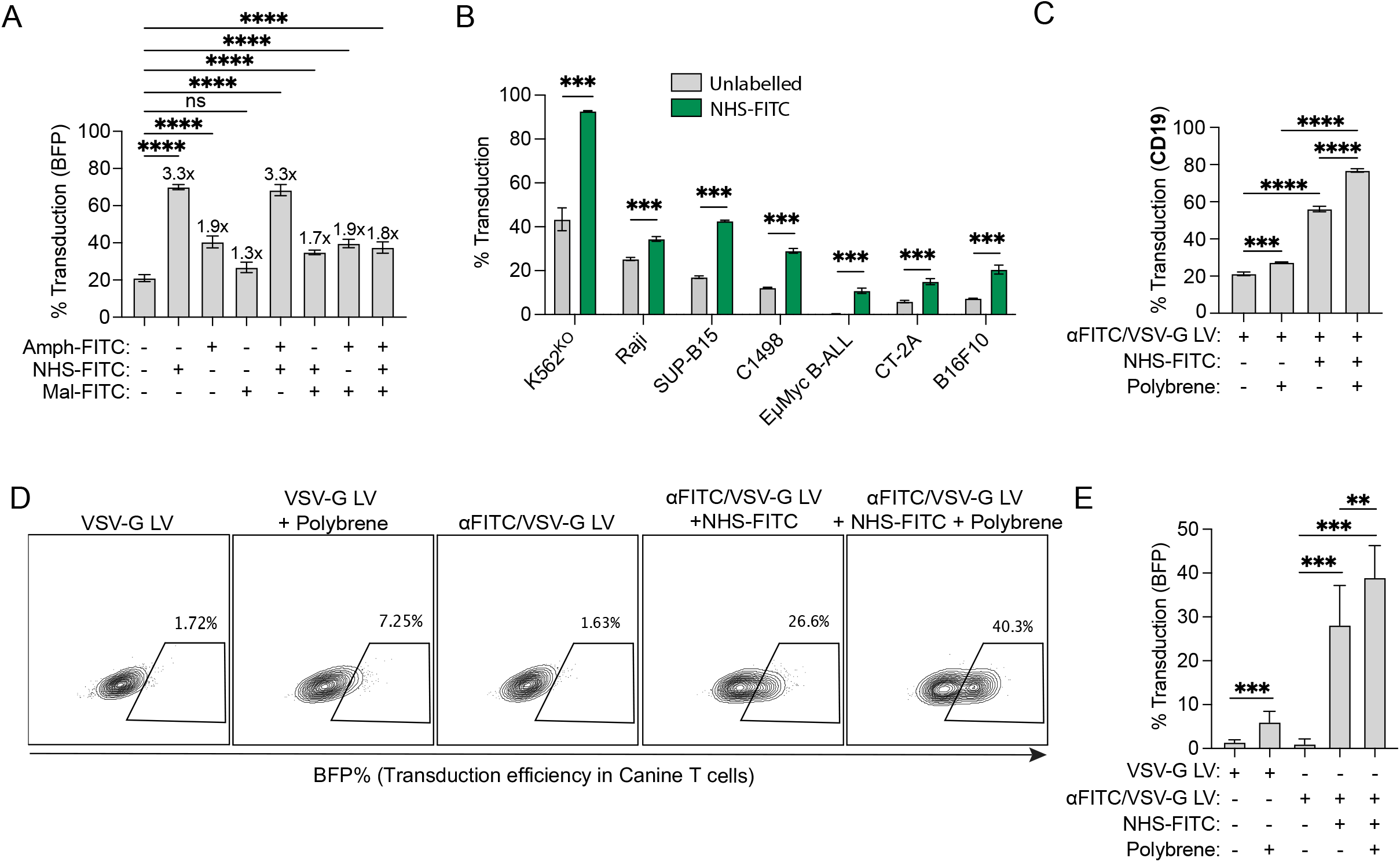
Paired cell surface FITC labeling and αFITC-Env mediated LV retargeting as a generic approach to improve LV transduction of established mammalian cell lines and primary canine T cells. **(A)** Impact of combination labeling on αFITC/VSV-G LV transduction. K562^KO^ cells were labeled with either NHS-FITC (1μM), amph-FITC^5k^ (1μM), Mal-FITC (5μM) or combination of these as indicated in the figure followed by transduction with αFITC/VSV-G LV. (**B)** αFITC/VSV-G LV transduction of various mammalian cell lines pre-labeled with NHS-FITC (1μM). **(C)** K562^KO^ cells were labeled with NHS-FITC (1μM) followed by transduction with αFITC/VSV-G LV delivering CD19 with or without polybrene (20μg/ml). The percentage of transduced (CD19^+^) K562^KO^ was measured using flow cytometry 48 hours post transduction. (**D-E)** αFITC/VSV-G LV transduction of activated canine T cells pre-labeled with NHS-FITC (1μM) in the presence or absence of polybrene (10ug/ml). Unlabeled canine T cells transduced with VSV-G LV with or without polybrene (10ug/ml) are included as control. Shown are representative plots (**D**) and percentage of transduced cells (**E**). BFP was used the reporter for A-B, D-E, and transduction efficiently was determined as the BFP level 48 hours post transduction. Error bars show mean ± SD (n=3). *, P<0.05; **, P<0.01; ***, P<0.001; ****, P<0.0001; ns, non-significant by unpaired t-test for B or one-way ANOVA with Dunnett’s post-test for A; with Tukey’s post-test for C and E.

### Surface engineering improved LV transduction of naïve human T cells

Recently, it was found that ex vivo-manufactured CAR T cells with memory-like and naïve phenotypes exhibited superior anti-tumor activities and long-term efficacy in vivo compared to CAR T cells manufactured with prior activation^28^. Importantly, at least 10x fewer memory-like/naïve CAR T cells are needed to achieve an equivalent therapeutic efficacy^29^. To assess whether our surface engineering approach could enhance LV transduction of naïve human T cells, we labeled fresh naïve human T cells with NHS-FITC and monitored αFITC/VSV-G LV transduction with a BFP reporter (**Fig. 5A**). We observed consistent transduction across multiple donors with a median of 20% transduction efficiency and more than 50% for some donors (**Fig. 5B-C**). And using this approach, 40% (without polybrene) to 70% (with polybrene) of naïve human T cells could be transduced with CD19-CAR-bearing LVs (**Fig. 5D**). Analysis of the T cell phenotypes confirmed that NHS-FITC assisted LV transduction does not significantly affect T cell differentiation, and most CD19-CAR-transduced cells maintained their naïve phenotype (**Fig. 5E-F**).

**Figure 5:**
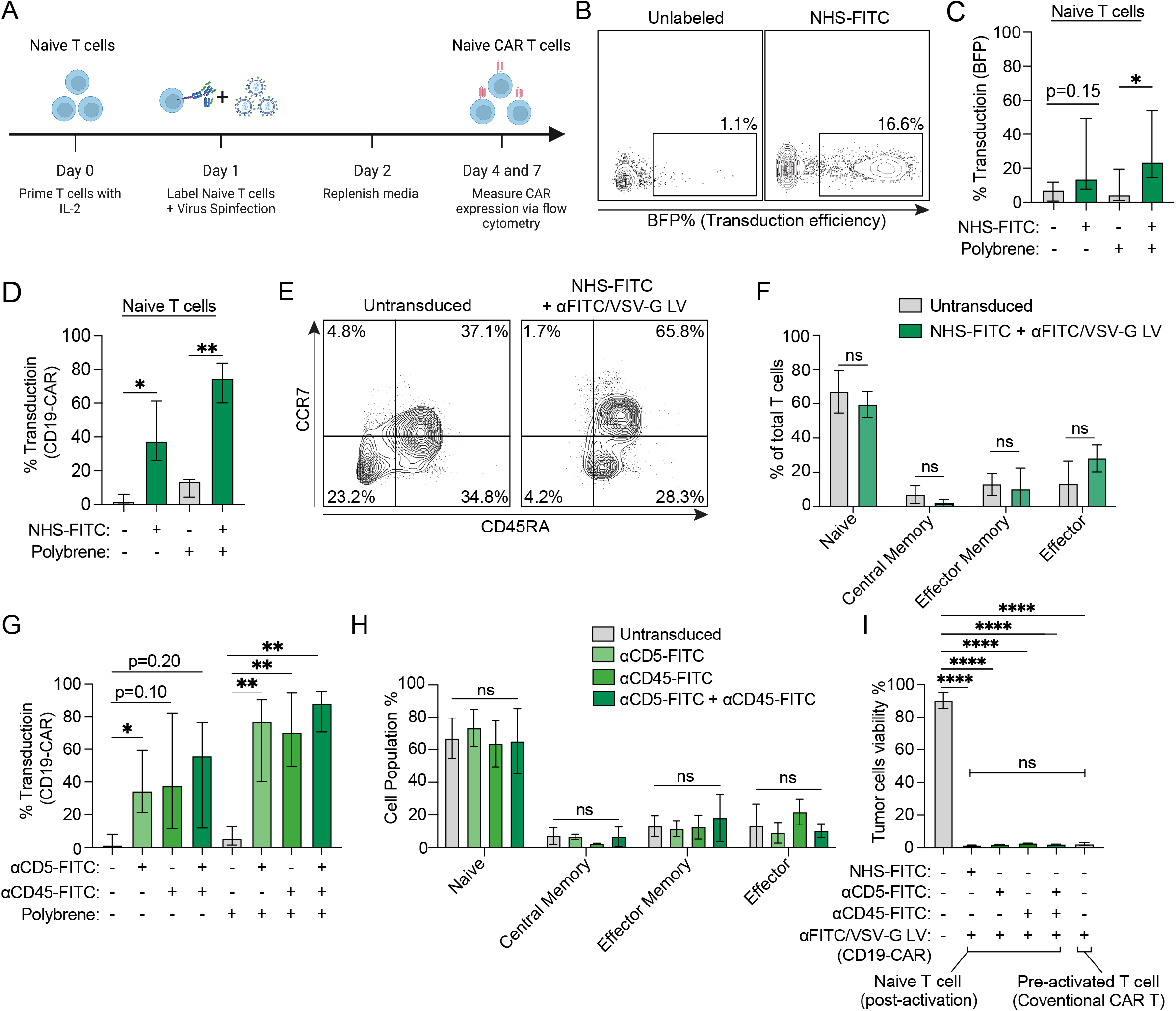
Cell surface engineering improved LV transduction of naïve human T cells. **(A)** Schematic timeline of naïve human T cell transduction. **(B-C)** αFITC/VSV-G LV (BFP) transduction of naïve human T cells (n=5) pre-labeled with or without NHS-FITC (1μM) and polybrene (5ug/ml). Shown are representative plots (**B**) and percentage of transduced cells (**C**). **(D)** αFITC/VSV-G LV (CD19-CAR) transduction of naïve human T cells (n=4) pre-labeled with or without NHS-FITC (1μM) and polybrene (5ug/ml). The percentage of transduced (CD19-CAR Flag tag^+^) naïve T cells was measured using flow cytometry 4 days post transduction. (**E-F)**. Immunophenotype of transduced (CD19-CAR Flag tag^+^) or untransduced (CD19-CAR Flag tag^-^) human T cells as in D. Shown are the representative plot (E) and the percentage (F) of naïve (CCR7^+^CD45RA^+^), central memory (CCR7^+^CD45RA^-^), effector memory (CCR7^-^CD45RA^-^), and effector cells (CCR7^-^CD45RA^+^). **(G)** FITC-antibody-directed αFITC/VSV-G LV (CD19-CAR) transduction of naïve human T cells. Naïve human T cells were labeled with either αCD5-FITC (n=6), αCD45-FITC (n=6), or a combination (n=4). The percentage of transduced (CD19-CAR Flag tag^+^) naïve T cells was measured using flow cytometry 4 days post transduction. **(H)** Immunophenotype of transduced (CD19-CAR Flag tag^+^) naive human T cells that are pre-labeled with various FITC-antibodies as in G. Untransduced control T cells (n=3) was used as control. Shown are the same untransduced samples as in Fig. 5E-F. **(I)** Cytotoxicity of naïve human CD19-CAR T cells after activation. Naïve human T cells were labeled with NHS-FITC, αCD5-FITC or αCD45-FITC followed by transduction with αFITC/VSV-G LVs (CD19-CAR). Two days after, T cells were activated for 5 days using αCD3/αCD28 dynabeads. T cell cytotoxicity was determined by co-culturing with luciferase-expressing CD19^+^ NALM6. Conventional pre-activated CD19-CAR T cells are included as control. Error bars show median ± 95% CI; *, P<0.05; **, P<0.01; ***, P<0.001; ****, P<0.0001; ns, by one way ANOVA with Šídák’s post-test for C; by mixed-effects analysis with Šídák post-test for D, G; by two-way ANOVA with Šídák’s post-test for F with Tukey’s post-test for H and by one way ANOVA with Dunnet’s post-test for I.

Considering that FITC-conjugated antibodies targeting various surface markers on T cells are commercially available, we sought to test if αFITC/VSV-G LV could transduce naïve human T cells pre-labeled with FITC-conjugated antibodies. To this end, naïve human T cells were pre-labeled with αCD5-FITC, αCD45-FITC or a combination of both (**Fig. 5G**), then transduced with αFITC/VSV-G LVs encoding the CD19-CAR. The median CD19 CAR transduction of naïve T cells increased from <5% to >30% (and >60% with the addition of polybrene), respectively (**Fig. 5G**). The transduced cells also maintained a naïve T cell phenotype (**Fig. 5H**). Lastly, these naïve CD19-CAR T cells could be activated and kill CD19+ Nalm6 cells efficiently in vitro, equivalent to CD19-CAR T cells manufactured using the conventional procedure **(Fig. 5I)**.

## DISCUSSION

Efficient and stable gene delivery to target cells entails LV recognition of a cell surface receptor that can facilitate virus docking and cell entry. Although VSV-G has been utilized for LV pseudotyping owing to its broad tropism, many cell types heavily used in pre-clinical research lack a high-level expression of LDLR family proteins that are required for VSV-G binding, resulting in limited or no gene delivery^30^. Although pseudotying LV with alternative Env protein could be leveraged to transduce certain cell types, it is generally required that the Env protein be well characterized for its target and the options of these alternative Env proteins are also limited^11^. Here we found that VSV-G and αFITC-Env dual pseudotying coupled with the synthetic addition of FITC to the cell surface enables enhanced LV gene delivery to a variety of cell types including cell types that are naturally resistant to VSV-G LV, such as canine T cells and naïve human T cells. This simple two-step approach for LV gene delivery has two major impacts: 1) provides a universal approach for efficient LV gene delivery into most mammalian cell types without the need for customized pseudotyping. 2) Applicable to both preclinical laboratory research and ex vivo manufacturing of therapeutic cells.

Mechanistically, dual VSV-G/αFITC-Env pseudotyping integrates the ability of VSV-G for efficient virus particle production and membrane fusion with αFITC-Env-mediated virus retargeting. Synthetic addition of FITC to the cell surface using chemical modification or antibody-based targeting endowed a generic docking ligand for LV on the surface of mammalian cells irrespective of the cells’ origin and genetic background, and cell surface FITC modification does not seem to affect cell viability. Notably, this generic approach enables virus to actively engage the target cell surface through a high-affinity FITC and αFITC-Env interaction in contrast to the passive virus attachment facilitated by cationic polymers such as polybrene. Although polybrene itself only has a mild effect, the addition of polybrene further promoted transduction in the presence of a proper entry receptor, likely through improving transient virus attachment to cell surface. These observations highlight that LV gene delivery is dictated by active virus binding to its receptor on the cell surface, which can be further enhanced by passive virus attachment.

One attractive feature of LV is that it can transduce non-dividing cells^31^. However, this property seems to be very inefficient for naïve human T cells. Recently, a highly sophisticated protocol was published to enable LV transduction of inactivated human T cells with a moderate efficiency of ∼25%, yet most of the transduced cells exhibited a central memory instead of naïve phenotype^32^. Using our FITC-labeling approach, we managed to transduce naïve human T cells with a markedly improved efficiency of 20-60%. Importantly, most of the transduced T cells exhibited a naïve phenotype. Post-activation of naïve CAR T cells resulted in fully functional CAR T cells. We recently developed a synthetic vaccine for boosting ex vivo-manufactured CAR T cells after adoptive transfer^27^. The booster vaccine will likely have a more pronounced effect on naïve CAR T cells than pre-activated T cells given their superior homing ability and differentiation potential. While this work is focused on in vitro assays for establishing and optimizing the enhanced LV gene delivery technology, we will evaluate the naïve CAR T generated using this approach in combination with vaccine boosting in vivo in a therapeutic setting in future studies.

In summary, we have developed a generic approach to enhance LV transduction of mammalian cells using paired cell surface FITC-labeling and VSV-G/αFITC-Env dual-pseudotyping. The non-selective FITC labeling makes it a near-universal and highly customizable solution to lentiviral gene delivery into mammalian cells. This approach has wide application for both routine lab research and therapeutic T-cell engineering.

## ACKNOWLEDGMENTS

We thank Dr. Michael Hemann from the Koch Institute of MIT for sharing the Eμ-Myc cells. We thank the Children’s Hospital of Philadelphia Research Vector Core for the HEK 293T cells. We thank the Birnbaum Lab at MIT for the 58^-/-^ Hybridoma cells. **Funding:** The work was supported by the Cell and Gene Therapy Collaborative and the Junior faculty pilot program at CHOP, NIH New Innovators Award (DP2 AI164319-03), ITMAT at UPenn and NCATS (UL1TR001878). A.N. is supported by a National Science Foundation Graduate Research Fellowship Program award.

## Data and materials availability

All data are available on request.

## AUTHOR CONTRIBUTIONS

R.T., L.C. and L.M. designed the experimental strategy. L.M. designed the engineered viral envelope. R.T., L.C., T.M.G., and L.M. wrote the manuscript. R.T., L.C., O.A., T.M.G. and S.K performed the experiments. N.J.M and L.O. prepared activated canine T cells. A.N. designed and produced the anti-VSV-G antibody. R.T., T.M.G., and L.M. analyzed and interpreted the data. R.T. and L.M. prepared the figures with the help of T.M.G.

## References

(1) Johnson, N. M.; Alvarado, A. F.; Moffatt, T. N.; Edavettal, J. M.; Swaminathan, T. A.; Braun, S. E. HIV-based lentiviral vectors: origin and sequence differences. Mol Ther Methods Clin Dev 2021, 21, 451–465. DOI: 10.1016/j.omtm.2021.03.018 From NLM.

(2) Breckpot, K.; Aerts, J. L.; Thielemans, K. Lentiviral vectors for cancer immunotherapy: transforming infectious particles into therapeutics. Gene Therapy 2007, 14 (11), 847–862. DOI: 10.1038/sj.gt.3302947.

(3) Irving, M.; Lanitis, E.; Migliorini, D.; Ivics, Z.; Guedan, S. Choosing the Right Tool for Genetic Engineering: Clinical Lessons from Chimeric Antigen Receptor-T Cells. Hum Gene Ther 2021, 32 (19-20), 1044–1058. DOI: 10.1089/hum.2021.173 From NLM.

(4) Cronin, J.; Zhang, X. Y.; Reiser, J. Altering the tropism of lentiviral vectors through pseudotyping. Curr Gene Ther 2005, 5 (4), 387–398. DOI: 10.2174/1566523054546224 From NLM.

(5) Lentz, T. L.; Burrage, T. G.; Smith, A. L.; Crick, J.; Tignor, G. H. Is the acetylcholine receptor a rabies virus receptor? Science 1982, 215 (4529), 182–184. DOI: 10.1126/science.7053569 From NLM.

(6) Bartz, R.; Firsching, R.; Rima, B.; ter Meulen, V.; Schneider-Schaulies, J. Differential receptor usage by measles virus strains. J Gen Virol 1998, 79 (Pt 5), 1015–1025. DOI: 10.1099/0022-1317-79-5-1015 From NLM.

(7) Witting, S. R.; Vallanda, P.; Gamble, A. L. Characterization of a third generation lentiviral vector pseudotyped with Nipah virus envelope proteins for endothelial cell transduction. Gene Ther 2013, 20 (10), 997–1005. DOI: 10.1038/gt.2013.23 From NLM.

(8) Hislop, J. N.; Islam, T. A.; Eleftheriadou, I.; Carpentier, D. C. J.; Trabalza, A.; Parkinson, M.; Schiavo, G.; Mazarakis, N. D. Rabies Virus Envelope Glycoprotein Targets Lentiviral Vectors to the Axonal Retrograde Pathway in Motor Neurons * <sup></sup>. Journal of Biological Chemistry 2014, 289 (23), 16148–16163. DOI: 10.1074/jbc.M114.549980 (acccessed 2024/09/25).

(9) Finkelshtein, D.; Werman, A.; Novick, D.; Barak, S.; Rubinstein, M. LDL receptor and its family members serve as the cellular receptors for vesicular stomatitis virus. Proc Natl Acad Sci U S A 2013, 110 (18), 7306–7311. DOI: 10.1073/pnas.1214441110 From NLM.

(10) Sun, X.; Yau, V. K.; Briggs, B. J.; Whittaker, G. R. Role of clathrin-mediated endocytosis during vesicular stomatitis virus entry into host cells. Virology 2005, 338 (1), 53–60. DOI: 10.1016/j.virol.2005.05.006 From NLM.

(11) Dautzenberg, I. J. C.; Rabelink, M. J. W. E.; Hoeben, R. C. The stability of envelope-pseudotyped lentiviral vectors. Gene Therapy 2021, 28 (1), 89–104. DOI: 10.1038/s41434-020-00193-y.

(12) Hanenberg, H.; Xiao, X. L.; Dilloo, D.; Hashino, K.; Kato, I.; Williams, D. A. Colocalization of retrovirus and target cells on specific fibronectin fragments increases genetic transduction of mammalian cells. Nat Med 1996, 2 (8), 876–882. DOI: 10.1038/nm0896-876 From NLM.

(13) Sutton, R. E.; Reitsma, M. J.; Uchida, N.; Brown, P. O. Transduction of human progenitor hematopoietic stem cells by human immunodeficiency virus type 1-based vectors is cell cycle dependent. J Virol 1999, 73 (5), 3649–3660. DOI: 10.1128/jvi.73.5.3649-3660.1999 From NLM.

(14) Kootstra, N. A.; Zwart, B. M.; Schuitemaker, H. Diminished human immunodeficiency virus type 1 reverse transcription and nuclear transport in primary macrophages arrested in early G(1) phase of the cell cycle. J Virol 2000, 74 (4), 1712–1717. DOI: 10.1128/jvi.74.4.1712-1717.2000 From NLM.

(15) Dardalhon, V.; Jaleco, S.; Kinet, S.; Herpers, B.; Steinberg, M.; Ferrand, C.; Froger, D.; Leveau, C.; Tiberghien, P.; Charneau, P.; et al. IL-7 differentially regulates cell cycle progression and HIV-1-based vector infection in neonatal and adult CD4+ T cells. Proc Natl Acad Sci U S A 2001, 98 (16), 9277–9282. DOI: 10.1073/pnas.161272698 From NLM.

(16) Serafini, M.; Naldini, L.; Introna, M. Molecular evidence of inefficient transduction of proliferating human B lymphocytes by VSV-pseudotyped HIV-1-derived lentivectors. Virology 2004, 325 (2), 413–424. DOI: 10.1016/j.virol.2004.04.038 From NLM.

(17) Kerkar, S. P.; Sanchez-Perez, L.; Yang, S.; Borman, Z. A.; Muranski, P.; Ji, Y.; Chinnasamy, D.; Kaiser, A. D.; Hinrichs, C. S.; Klebanoff, C. A.; et al. Genetic engineering of murine CD8+ and CD4+ T cells for preclinical adoptive immunotherapy studies. J Immunother 2011, 34 (4), 343–352. DOI: 10.1097/CJI.0b013e3182187600 From NLM.

(18) Panjwani, M. K.; Atherton, M. J.; MaloneyHuss, M. A.; Haran, K. P.; Xiong, A.; Gupta, M.; Kulikovsaya, I.; Lacey, S. F.; Mason, N. J. Establishing a model system for evaluating CAR T cell therapy using dogs with spontaneous diffuse large B cell lymphoma. Oncoimmunology 2020, 9 (1), 1676615. DOI: 10.1080/2162402x.2019.1676615 From NLM.

(19) Amirache, F.; Lévy, C.; Costa, C.; Mangeot, P.-E.; Torbett, B. E.; Wang, C. X.; Nègre, D.; Cosset, F.-L.; Verhoeyen, E. Mystery solved: VSV-G-LVs do not allow efficient gene transfer into unstimulated T cells, B cells, and HSCs because they lack the LDL receptor. Blood 2014, 123 (9), 1422–1424. DOI: 10.1182/blood-2013-11-540641 (acccessed 9/26/2024).

(20) Bukrinsky, M. I.; Sharova, N.; Dempsey, M. P.; Stanwick, T. L.; Bukrinskaya, A. G.; Haggerty, S.; Stevenson, M. Active nuclear import of human immunodeficiency virus type 1 preintegration complexes. Proc Natl Acad Sci U S A 1992, 89 (14), 6580–6584. DOI: 10.1073/pnas.89.14.6580 From NLM.

(21) Vatakis, D. N.; Kim, S.; Kim, N.; Chow, S. A.; Zack, J. A. Human immunodeficiency virus integration efficiency and site selection in quiescent CD4+ T cells. J Virol 2009, 83 (12), 6222–6233. DOI: 10.1128/jvi.00356-09 From NLM.

(22) Stevenson, M.; Stanwick, T. L.; Dempsey, M. P.; Lamonica, C. A. HIV-1 replication is controlled at the level of T cell activation and proviral integration. Embo j 1990, 9 (5), 1551–1560. DOI: 10.1002/j.1460-2075.1990.tb08274.x From NLM.

(23) Fichter, C.; Aggarwal, A.; Wong, A. K. H.; McAllery, S.; Mathivanan, V.; Hao, B.; MacRae, H.; Churchill, M. J.; Gorry, P. R.; Roche, M.; et al. Modular Lentiviral Vectors for Highly Efficient Transgene Expression in Resting Immune Cells. Viruses 2021, 13 (6). DOI: 10.3390/v13061170 From NLM.

(24) Agarwal, S.; Hanauer, J. D. S.; Frank, A. M.; Riechert, V.; Thalheimer, F. B.; Buchholz, C. J. In Vivo Generation of CAR T Cells Selectively in Human CD4(+) Lymphocytes. Mol Ther 2020, 28 (8), 1783–1794. DOI: 10.1016/j.ymthe.2020.05.005 From NLM. Pfeiffer, A.; Thalheimer, F. B.; Hartmann, S.; Frank, A. M.; Bender, R. R.; Danisch, S.; Costa, C.; Wels, W. S.; Modlich, U.; Stripecke, R.; et al. <i>In vivo</i> generation of human CD19‐CAR T cells results in B‐cell depletion and signs of cytokine release syndrome. EMBO Molecular Medicine 2018, 10 (11), e9158. DOI: 10.15252/emmm.201809158 (acccessed 2024/09/03). Dreja, H.; Piechaczyk, M. The effects of N-terminal insertion into VSV-G of an scFv peptide. Virology Journal 2006, 3 (1), 69. DOI: 10.1186/1743-422X-3-69.

(25) O’Doherty, U.; Swiggard, W. J.; Malim, M. H. Human immunodeficiency virus type 1 spinoculation enhances infection through virus binding. J Virol 2000, 74 (21), 10074–10080. DOI: 10.1128/jvi.74.21.10074-10080.2000 From NLM.

(26) Davis, H. E.; Morgan, J. R.; Yarmush, M. L. Polybrene increases retrovirus gene transfer efficiency by enhancing receptor-independent virus adsorption on target cell membranes. Biophys Chem 2002, 97 (2-3), 159–172. DOI: 10.1016/s0301-4622(02)00057-1 From NLM.

(27) Ma, L.; Dichwalkar, T.; Chang, J. Y. H.; Cossette, B.; Garafola, D.; Zhang, A. Q.; Fichter, M.; Wang, C.; Liang, S.; Silva, M.; et al. Enhanced CAR-T cell activity against solid tumors by vaccine boosting through the chimeric receptor. Science 2019, 365 (6449), 162–168. DOI: 10.1126/science.aav8692 From NLM.

(28) López-Cantillo, G.; Urueña, C.; Camacho, B. A.; Ramírez-Segura, C. CAR-T Cell Performance: How to Improve Their Persistence? Front Immunol 2022, 13, 878209. DOI: 10.3389/fimmu.2022.878209 From NLM.

(29) Arcangeli, S.; Bove, C.; Mezzanotte, C.; Camisa, B.; Falcone, L.; Manfredi, F.; Bezzecchi, E.; El Khoury, R.; Norata, R.; Sanvito, F.; et al. CAR T cell manufacturing from naive/stem memory T lymphocytes enhances antitumor responses while curtailing cytokine release syndrome. J Clin Invest 2022, 132 (12). DOI: 10.1172/jci150807 From NLM.

(30) Jin, H.; Zhang, C.; Zwahlen, M.; von Feilitzen, K.; Karlsson, M.; Shi, M.; Yuan, M.; Song, X.; Li, X.; Yang, H.; et al. Systematic transcriptional analysis of human cell lines for gene expression landscape and tumor representation. Nature Communications 2023, 14 (1), 5417. DOI: 10.1038/s41467-023-41132-w.

(31) Mátrai, J.; Chuah, M. K. L.; VandenDriessche, T. Recent Advances in Lentiviral Vector Development and Applications. Molecular Therapy 2010, 18 (3), 477–490. DOI: 10.1038/mt.2009.319 (acccessed 2024/10/24).

(32) Ghassemi, S.; Durgin, J. S.; Nunez-Cruz, S.; Patel, J.; Leferovich, J.; Pinzone, M.; Shen, F.; Cummins, K. D.; Plesa, G.; Cantu, V. A.; et al. Rapid manufacturing of non-activated potent CAR T cells. Nature Biomedical Engineering 2022, 6 (2), 118–128. DOI: 10.1038/s41551-021-00842-6.

(33) Ma, L.; Boucher, J. I.; Paulsen, J.; Matuszewski, S.; Eide, C. A.; Ou, J.; Eickelberg, G.; Press, R. D.; Zhu, L. J.; Druker, B. J.; et al. CRISPR-Cas9-mediated saturated mutagenesis screen predicts clinical drug resistance with improved accuracy. Proc Natl Acad Sci U S A 2017, 114 (44), 11751–11756. DOI: 10.1073/pnas.1708268114 From NLM.

