## Supplementary Material for "Enhanced lentiviral gene delivery to mammalian cells via paired cell surface and viral envelope engineering"

### MATERIALS AND METHODS

#### Cell Culture

K562 (CCL-243), Jurkat (TIB-152), Nalm6 (CRL-3273) were obtained from ATCC and cultured in RPMI 1640 medium (Corning). HEK293T cells were a gift from the Children's Hospital of Philadelphia Research Vector Core. HEK293T cells were maintained in High-Glucose DMEM medium (Corning). C1498 (TIB-49) were obtained from ATCC and cultured in High-Glucose DMEM medium (Corning). SUP-B15 were obtained from ATCC (CRL-1929) and cultured in IMDM (Gibco) supplemented with 4 mM L-glutamine. 58-/- Hybridoma were a gift from the Birnbaum lab at MIT. All cell lines were cultured in media supplemented with 10% Fetal Bovine Serum (HyClone) and 100 units/mL of penicillin and 100 µg/mL of streptomycin (Gibco). ExpiCHO cells were obtained from ThermoFisher and cultured in ExpiCho Expression Medium (Gibco). The Eµ-Myc cell line was a gift from Dr. Michael Hemann at MIT. Cells were cultured in a medium composed of a 50:50 mix of IMDM with L-glutamine and 25 mM HEPES (Gibco) and DMEM with L-glutamine and sodium pyruvate (Corning), supplemented with 10% FBS and 2-mercaptoethanol to a final concentration of 0.05 mM (Gibco). CT2A (SCC194) and B16F10 (CRL-6475) cells were obtained from ATCC and cultured in High-Glucose DMEM medium (Corning).

#### Generation of LDLR-KO K562 cells

The CRISPR Cas9 cassette targeting the hLDLR gene locus (sgRNA target site: GCTGCGAGCATGGGGCCCTG) and containing puromycin resistance gene was introduced into K562 cells via transduction with lentiviral vectors. Transduced cells were selected using puromycin at 1 µg/mL for 6 days. LDLR expression was measured via flow cytometry.

#### Antibodies and Recombinant Proteins

The heavy chain and light chain of the anti-VSV-G IgG (clone: I1) was amplified and sequenced as previously described[33]. Then each chain was cloned into the gWiz vector and combined during transfection for recombinant protein production using ExpiCho cells. ExpiCho cells were seeded in ExpiCho Expression Medium (Gibco) at a density of $3x10^6$ cells/ml. After 24 hours, cells were transfected with 1:3 ratio of heavy to light chain plasmids according to the manufacturer's protocol using ExpiFectamine CHO Transfection Kit (Gibco, Cat#: A29130). Antibody supernatant was collected 10 days after transfection and filtered using a 0.45 µm filter (Globescientific, Cat#: SF-PVDF-4513-S). Recombinant proteins were then purified using FPLC (Akta pure 25). Anti-human CD19 PE (Clone: HIB19), anti-CD71 FITC (Clone: CY1G4), anti-CD5 FITC (Clone: UCHT2), anti-CD45-FITC (Clone: 30-F11), anti-DYKDDDDK Tag APC (Clone: anti-Flag Tag), anti-CCR7 PE (Clone: G043H7) and anti-CD45RA Alexa Fluor 488 (Clone: HI100) antibodies were all purchased from Biolegend.

#### Construction of lentiviral vectors

The human CD19-CAR construct was generated by fusing geneblock fragments (custom ordered from IDT) into a lentiviral vector containing the EF1α promoter. The CD19-CAR is composed of a human CD8 signal peptide, FMC63 scFv (VLVH), human CD8 hinge and transmembrane domain, 41BB costimulatory domain and CD3ζ intracellular domain. To facilitate CAR detection by flow cytometry, a flag tag was inserted at the N-terminus of FMC63 scFv immediately following the signal peptide. The human CD19 construct was generated by fusing geneblock fragments (custom ordered from IDT) into a lentiviral vector containing the EF1α promoter.

#### Lentiviral Production

For lentivirus production, HEK293T cells were cultured till 70% confluency, then split at 1:3 for further expansion. 24 hours later, HEK293T cells were seeded at $3.75x10^6$ cells/dish in 10 cm tissue culture dishes and cultured for 16 hours till the confluency reached 70-80%. Complete media was replenished after 16 hours and cells were transfected with polyethyleneimine (PEI) 25k (Polysciences Cat#: 23966-1). Briefly, for each transfection, 10 µg DNA in a ratio of 4:1:1:4 of psPAX2, pMD2.G, αFITC-Env, and a plasmid encoding the gene of interest (either BFP, human CD19, or CD19 CAR), were mixed with 1 ml of Opti-MEM (Gibco), and 30 µL of 1 mg/mL PEI in a ratio of 3:1 PEI to DNA. Mixture was incubated at room temperature for 20 minutes before being added to the 10 cm dishes of HEK293T cells. For VSV-G lentivirus, a ratio of 3:3:4 of psPAX2, pMD2.G and a plasmid encoding gene of interest, was used. After 24 hours, media was replenished. The lentiviral supernatant was collected 48 hours after media replenishment and clarified by centrifugation at 2,000xg for 10 mins and stored at -80°C until further use.

**Lentiviral Transduction**

Target cells were seeded into a 48-well plate tissue culture plate at a density of $1x10^5$ cells in 200 µL volume of media with or without 20 µg/ml of polybrene. Virus-containing supernatant was added, and the cells were spun for 1.5 hours at 2000xg and 35°C. 100 µl of complete media was added to each well after transduction. Transduction efficiency was measured via flow cytometry at the day 2 or day 7 as indicated in the figure. For some experiments, cells were transduced with a retronectin plate coated with 15 µg/ml (Takara, Cat#: T100B) and spun at 2000xg for 1.5 hours.

**Chemical Labeling**

Cells were washed with 1xDPBS (Corning) and suspended at a density of $1x10^6$ cells/ml in 1xDPBS with 5-Carboxyfluorescein Succinimidyl Ester (NHS-FITC, ThermoFischer, Cat#: C2210), DSPE-PEG-FITC MW, 1k, 2k, 3.4k, 5k, or 10k (Amph-FITC, Creative PEG Works, Cat#: PLS-9926, PLS-9927, PLS-9928, PLS-9929, PSB-2253) or Fluorescein 5 Maleimide (Maleimide-FITC, ThermoFisher, Cat#: 62245) with concentration as indicated in the figures. Cells were incubated with the chemical for 40 minutes at 37°C. Following the incubation, cells were washed with complete media. To determine the accessibility of the FITC at the cell membrane, cells were stained with Alexa Fluor 647 anti-FITC antibody (Jackson ImmunoResearch, Cat#: 200-602-037) at a concentration of 1 µg/ml for 20 minutes on ice, followed by analysis on BD LSRII.

**Two-step Click Chemistry**

Cells were washed with 1xDPBS (Corning) and suspended at a density of $1x10^6$ cells/ml in 1xDPBS and labeled with 1 µM DBCO-NHS Ester (NHS-DBCO, ClickChemistryTools, Cat#: A133-25) or 40 minutes at 37°C. Cells were washed with complete media and incubated with 1 µM 5-FAM azide (FITC-azide, Lumiprobe, Cat#: C4130) for 40 minutes at 37°C and cells were shaken every 10 minutes. Cells were subsequently washed with complete media and used for transduction.

**Lentivirus Binding**

Following chemical or antibody labeling, $1x10^5$ cells were spun with VSV-G-LV or αFITC/VSV-G LV in complete media for 10 minutes at 2000xg. Cells were incubated at 37°C for 20 minutes, following spin. Unbound virus was washed with FACS buffer (DPBS containing 1 mM EDTA (Sigma), 0.5% Bovine Serum Albumin (Sigma), and 0.1% Sodium azide (Sigma)). Cells were stained with anti-VSV-G antibody on ice at a concentration of 1 µg/ml for 20 minutes. Cells were washed FACS buffer and stained for 20 minutes on ice with an anti-murine IgG2a PE (Biolegend, Clone: RMG2a-62) antibody at a final concentration of 1 µg/ml. Cells were then washed with FACS buffer and resuspended for analysis via flow cytometry on BD LSRII.

**Blocking VSV-G Binding**

VSV-G binding to LDLR was blocked by incubating lentivirus-containing supernatant with an anti-VSV-G antibody at a concentration of 1 µg/ml for 20 minutes at room temperature. Pre-blocked virus was subsequently used in certain groups in binding and transduction experiments as indicated.

**Flow Cytometry and Analysis**

Cells were prepared for flow cytometry by washing with FACS buffer (1xDPBS containing 1 mM EDTA, 0.5% Bovine Serum Albumin, and 0.1% Sodium azide) and re-suspending in 7AAD live/dead dye (Biolegend, Cat#: 420404) or Near-IR live dead reactive dye (Thermofisher Scientific, Cat#: L10119) according to the manufacture protocol. Cells were washed with FACS buffer and resuspended at a density of $1x10^6$ cells per 100 µl in FACS buffer, followed by labeling with antibodies at 1 µg/ml for 20 minutes on ice. Cells were then washed with FACS buffer and fluorescence level was measured using the BD LSRII, BD Fortessa X-20 or CytoFLEX LX cytometer. Cell debris was excluded followed by exclusion of doublets and subsequently gated on live cells. Gating of positive population was adjusted based on the appropriate controls, as indicated in the figures. FlowJo 10.10 software was used to analyze the data.

**Naïve T cell Transduction**

$CD3^+$ T cells were isolated by negative selection using RosettaSep Kits from STEMCELL Technologies and obtained from Human Immunology Core at the Perelman School of Medicine at the University of Pennsylvania. Cells were cultured with human interleukin-2 (hIL-2) (PeproTech, Cat#: 200-02-50UG) at a concentration of 100 IU/ml for 24 hours in complete media (RPMI 1640 containing 10% FBS and 100 units/mL of penicillin and 100 µg/mL of streptomycin), at 37°C. Cell labeling and antibody labeling with anti-CD5-FITC (Biolegend, Clone: UCHT2), and anti-CD45-FITC (Biolegend, Clone: 30-F11) were performed as described in the flow analysis method. $1x10^6$ cells were spun in a 24-well tissue culture plate at with αFITC/VSV-G LV with or without 5 µg/ml polybrene at 2000xg for 1.5 hours at 35°C. After the spin, cells were transferred to the incubator and complete media containing IL-2 was refreshed after 24 hours. Transduction efficiency was determined using flow cytometry on day 2 post transduction, day 4 post transduction, or day 7 post transduction, as indicated in the figures.

**Cell Proliferation and Viability Assay**

Target cells were seeded into a 48-well plate tissue culture plate at a density of $1x10^5$ cells in 200 µL volume of media, chemically labeled and transduced as described before. Live cells were counted on the indicated days, using a light microscope and trypan blue staining.

**Conventional CAR T manufacturing Protocol**

$CD3^+$ T cells were isolated by negative selection using RosettaSep Kits from STEMCELL Technologies and obtained from Human Immunology Core at the Perelman School of Medicine at the University of Pennsylvania. T cells were activated with Human T-Activator CD3/CD28 Dynabeads (ThermoFisher, Cat#: 11131D) at a bead-to-cell ratio of 1:1 in complete medium supplemented with 100 IU/mL recombinant human IL-2 (PeproTech, Cat#: 200-02-50UG). After 24 hours of activation, T cells were transduced with human CD19 CAR lentiviral-supernatant with 5 µg/ml polybrene and spinfection was performed at 2000xg for 90 minutes at 35°C. Plates were then carefully transferred to an incubator and maintained overnight. The next day, plates were centrifuged at 600xg for 5 minutes and lentivirus-containing supernatant was removed and replaced with new virus. Spinfection was repeated as previously described. Cells were transferred to the incubator and after 8 hours the cell media was replenished with new complete media containing 100 IU/mL recombinant human IL-2. Transduction efficiencies were determined by flow cytometry 2 days later using anti-flag antibody.

**CAR T cell Cytotoxicity Assay**

Naïve human CAR T cells expressing CD19 CAR were activated with Human T-Activator CD3/CD28 Dynabeads (ThermoFisher, Cat#: 11132D) at a bead-to-cell ratio of 1:1 in complete medium supplemented with 100 IU/mL recombinant human IL-2 (PeproTech, Cat#: 200-02-50UG) for 5 days. CAR+ T cells were enriched by staining total expanded T cells with a PE-conjugated anti-flag antibody followed by staining with anti-PE microbeads and magnetic selection for CAR+ T cells according to the manufacturer's protocol (Miltenyi). Enriched CAR+ cells were co-cultured with luciferase-expressing CD19+ Nalm6 cells for 20 hours at an effector to target (E:T) ratio of 3:1. D-luciferin substrate (Perkin Elmer, Cat#: 122799) was added at a concentration of 150 µg/ml and bioluminescence was measuring using a plate reader (Tecan). Luciferase activity was normalized to targets only and presented as tumor cell viability percentage.

**Statistical Analysis**

Statistical analyses were performed using GraphPad Prism 10. All pair-wise comparisons were analyzed by student's t-test. Multi-group comparisons were carried out using a one-way ANOVA with Tukey's post-test, Dunnett's post-test, or Šidák's post-test as indicated in the figure legends , curve fitted and $R^2$ calculated by simple linear regression, by two-way ANOVA with Tukey's post-test, or by mixed-effects analysis with Šídák post-test or by two-way ANOVA with Šídák's post-test. Data are shown as means ± SD or median ± 95% CI, as indicated in the figure legends. Methods of statistical analyses are defined in every figure legend. A P value of less than 0.05 was statistically significant. Each experiment was performed in technical triplicates.
